# A bispecific antibody NXT007 exerts a hemostatic activity in hemophilia A monkeys enough to keep a non-hemophiliac state

**DOI:** 10.1101/2022.12.19.520692

**Authors:** Yuri Teranishi-Ikawa, Tetsuhiro Soeda, Hikaru Koga, Kazuki Yamaguchi, Kazuki Kato, Keiko Esaki, Kentaro Asanuma, Miho Funaki, Mina Ichiki, Yuri Ikuta, Shunsuke Ito, Eri Joyashiki, Shun-Ichiro Komatsu, Atsushi Muto, Kei Nishimura, Momoko Okuda, Hisakazu Sanada, Motohiko Sato, Norihito Shibahara, Tetsuya Wakabayashi, Koji Yamaguchi, Akiko Matsusaki, Zenjiro Sampei, Hirotake Shiraiwa, Hiroko Konishi, Yoshiki Kawabe, Kunihiro Hattori, Takehisa Kitazawa, Tomoyuki Igawa

## Abstract

Emicizumab, a factor (F)VIIIa-function mimetic bispecific antibody (BsAb) to FIXa and FX, has become an indispensable treatment for people with hemophilia A (PwHA). Although emicizumab is very potent, long-term outcomes from the clinical studies suggest that a small proportion of PwHA still experiences bleeds. Additionally, non-clinical studies indicate that the maximum cofactor activity of emicizumab is lower than international standard activity (100 IU/dL of FVIII). An increased cofactor activity BsAb would benefit such patients. Here, we report NXT007, a BsAb binding FIXa and FX developed through further engineering of emicizumab. Emicizumab has a common light chain, but through advances in antibody engineering, we were able to create a more potent BsAb with two new non-common light chains. After extensive optimization of the heavy and light chains, the resulting BsAb, NXT007, exerted in vitro thrombin generation (TG) activity in hemophilia A plasma equivalent to 100 IU/dL of FVIII when triggered by tissue factor. NXT007 demonstrated potent hemostatic activity in an acquired hemophilia A model in non-human primates at a much lower dosage than emicizumab, consistent with an around 30-fold dose shift in the in vitro TG activity between NXT007 and emicizumab. Moreover, together with Fc engineering that enhanced FcRn binding and reduced in vivo clearance, we demonstrate that NXT007 could be effective at a much lower dosage with a longer dosing interval compared to emicizumab. These non-clinical results suggest that NXT007 could maintain a non-hemophilic range of coagulation potential in PwHA and provides a rationale for its clinical testing.

## INTRODUCTION

Congenital hemophilia A, which is a disease caused by the deficiency or malfunction of factor VIII (FVIII), is classified into three main forms: severe, moderate, and mild.^1^ About half of people with hemophilia A (PwHA) are classified with severe disease, which is defined as having less than 1 IU/dL of FVIII activity in their plasma (i.e., <1% of the standard normal plasma).^2^ Moderate and mild diseases show 1–5 IU/dL and 5–40 IU/dL of the activity, respectively.^1,3^ PwHA with severe disease are at an especially high risk of bleeding, and would typically suffer frequent spontaneous bleedings into their joints and muscles as well as risk of intracranial bleeding without an appropriate bleeding-prophylactic treatment.^1^

Prophylactic treatment for PwHA has been established by regular infusions of a FVIII agent, which was proven to effectively prevent the majority of clinical bleeds, albeit with residual concerns about subclinical bleeds and long-term joint damage.^4-6^ However, FVIII-prophylactic treatments still typically require two or three weekly intravenous administrations due to its relatively short half-life,^7,8^ which imposes a substantial treatment burden. Although extended half-life products have been launched or are in development to reduce dosing frequency, the administration route has remained intravenous.^7^ Subcutaneous administration of FVIII has not been possible due to bioavailability and immunogenicity challenges.^9^ More critically, a remaining challenge is that one third of severe PwHA develop anti-FVIII alloantibodies, FVIII-inhibitors, which render FVIII agents ineffective.^10^

Emicizumab, launched in 2017, overcomes many of the shortcomings of FVIII agents. It is a therapeutic bispecific antibody (BsAb) to activated factor IX (FIXa) and factor X (FX) which mimics the cofactor activity of activated FVIII (FVIIIa).^11,12^ Owing to its long half-life and high bioavailability after subcutaneous administration, emicizumab became the first subcutaneously injectable agent for bleeding-prophylaxis. It achieved steady higher cofactor activity with a markedly lower dosing frequency (once a week, once every 2 weeks or once every 4 weeks) when compared to FVIII agents, making it much less burdensome, with a favorable reduction in treated bleeds compared to FVIII prohylaxis.^13^ Longer-term follow up study suggests >80% of patients achieved freedom from treated bleeds over prolonged periods of time.^14^ Moreover, emicizumab is effective even for PwHA who have FVIII-inhibitors^15-17^ because it is not recognized by FVIII-inhibitors due to its different structure from FVIII.

However, in vitro study and in vivo experience suggest emicizumab cofactor activity remains in the mild hemophilia range of FVIII:C equivalence.^15,18^ Thus, there was still room to increase the cofactor activity of a FVIIIa-mimetic BsAb, and that higher cofactor activity would be needed for those PwHA who continue to experience bleeding episodes while receiving emicizumab or clotting factor-based prophylaxis.

In this study, we aimed to create a new FVIIIa-function mimetic BsAb with higher cofactor activity, targeting the levels of 100 IU/dL of FVIII (equivalent to standard normal plasma) in vitro. By enhancing the activity, non-hemophiliac ranges of coagulation (40–150 IU/dL of FVIII-equivalence) could be constantly maintained in PwHA with a convenient dosing regimen. Given that the binding epitopes in FIX(a) and FX of emicizumab have been clinically validated to be effective and safe in PwHA, we decided to further engineer emicizumab by using the heavy chains as templates since they are the main contributors to the antigen binding.^19^ Moreover, we believed that use of the common light chain in emicizumab, which had been necessary at the time of its discovery,^12^ limited the potential for antibody optimization. Therefore, we decided to identify new pairs of light chains for each emicizumab heavy chain, then introduce mutations onto both heavy and light chains to enhance cofactor activity. After intensive screening, we identified a new BsAb, named NXT007. Non-clinical studies of NXT007 suggested that it has potential to maintain a non-hemophilic range of coagulation potential in PwHA justifying further clinical development.

## METHODS

### Identification of novel non-common light chains from phage library

The supplementary methods provide a detailed description of the identification of new light chains using the light chain shuffled Fab-displayed phage libraries.

### Expression and purification of bispecific antibodies

To generate a series of asymmetric BsAb variants consisting of two heavy chains and two light chains, each anti-FIXa and anti-FX antibody was produced separately by the Expi293 expression system (ThermoFisher Scientific) then combined for screening. Proprietary electrostatic steering mutations were introduced into the C_H_3 region of each heavy chain to promote heterodimerization.^20^ In producing NXT007, electrically charged mutation pairs were also introduced at each C_H_1/C_L_ interface to form the correct pairing of heavy chain and light chain.^21^ Each gene was inserted into the Expi293 expression system followed by purification using protein A and cation ion exchange chromatography to remove mis-paired moieties. After purification, the BsAb ratio was approximately 95%.

### Binding kinetics analysis by surface plasmon resonance (SPR)

The binding kinetics of NXT007 to human factor IX (hFIX), activated human factor IX (hFIXa), human factor X (hFX), activated human factor X (hFXa), cynomolgus monkey factor IX (cyFIX), or cynomolgus monkey factor X (cyFX) were assessed at pH 7.4 at 25°C using a Biacore T200 instrument (Cytiva). Detailed assay descriptions are provided in the supplementary methods.

### Enzymatic assay for FIXa-catalyzed FX activation

We evaluated the conversion rate of FX to FXa in an enzymatic assay using purified coagulation factors as described previously^11^ with a slight modification of phospholipid concentration and incubation time. Detailed assay descriptions are provided in the supplemental methods.

### Thrombin generation (TG) assays

Thrombin generation (TG) in plasma sample was measured by Calibrated Automated Thrombography (CAT, Thrombinoscope BV) using a 96-well plate fluorometer (ThermoFisher Scientific) as described previously.^18,22,23^ Briefly, each well in a 96-well plate was dispensed with 80 µL of plasma sample, to which was then added 20 µL of FXIa triggering solution consisting of 31 pM human FXIa (Enzyme Research Laboratories) and 20 µM synthetic phospholipid (10% phosphatidylserine, 60% phosphatidyl-choline, and 30% phosphatidylethanolamine), or TF triggering solution using PPP-reagent low (Thrombinoscope BV). Detailed assay descriptions for TG assays in cynomolgus monkey plasma are provided in the supplementary methods.

### Activated partial thromboplastin time (APTT) and prothrombin time (PT)

We employed as Thrombocheck APTT-SLA (Sysmex) for the APTT reagent and RecombiPlasTin 2G (Instrumentation Laboratory) for PT reagent using CS-2000i (Sysmex) for measurement as the recommended protocol. Detailed assay descriptions are provided in the supplementary methods.

### The care and use of laboratory animals

We used 37 male cynomolgus monkeys for the in vivo hemostatic study (2.6–4.0 kg, 3–4 years; Shin Nippon Biomedical Laboratories, Ltd., Kagoshima, Japan), 16 males for the pharmacokinetic study (3.0–3.8 kg, 3 years; Shin Nippon Biomedical Laboratories, Ltd., Kagoshima, Japan) and 20 females and 20 males for the toxicology study (6.0–9.5 kg, 6–11 year old males and 2.8–5.9 kg, 4–9 year old females; Shin Nippon Biomedical Laboratories, Ltd., Kagoshima, Japan). All animal studies were approved by the Institutional Animal Care and Use Committee of Chugai Pharmaceutical Co., Ltd., and were conducted in accordance with the approved protocols and the Guidelines for the Care and Use of Laboratory Animals at the company. Shin Nippon Biomedical Laboratories and Chugai Pharmaceutical Co., Ltd. are fully accredited by The Association for Assessment and Accreditation of Laboratory Animal Care International (AAALAC).

### In vivo hemostatic study in an acquired hemophilia A model

Since NXT007 is highly species specific in its FVIIIa-mimetic cofactor activity similar to its template, emicizumab, in vivo hemostatic activity was evaluated using the non-human primate bleeding model as previously described^18^ with slight modification. Briefly, an acquired hemophilia A state was induced by the administration of anti-FVIII neutralizing antibody, cyVIII-2236. Bleeding was induced by muscle needling and subcutaneous exfoliation at abdomen, and then NXT007 was intravenously administered at a range of doses 0.0075 to 0.6 mg/kg, with measured bruised areas and decrease in blood hemoglobin level at specified timepoints. During the experimental period, one animal in the vehicle group encountered lethal bleeding at Day 2 possibly due to the intensity of the bleeding induction method. The animal was excluded from the analysis. Additional detailed assay descriptions are provided in the supplementary methods. The plasma NXT007 concentration was determined by the method described in the Pharmacokinetic study and simulation section.

### Pharmacokinetic study and simulation

NXT007 was administered intravenously (2□mg/kg;□*n*□=□4) or subcutaneously (0.02, 0.2 or 2 mg/kg;□*n*□=□4 for each group) to cynomolgus monkeys. For animals dosed intravenously, blood was sampled with a heparinized syringe at 5 min, 2, 8, 24, 48, 72, 96□hours, 7, 14, 21, 28, 35, 42, 49, 56, 63, 70, 77, and 84□days post dose. For animals dosed subcutaneously, blood was collected in the same way without sampling at 5□min. The collected blood was immediately centrifuged to separate the plasma, which was kept at −80°C until measurement. Details on the measurement of the concentration of NXT007 and data analysis are provided in the supplementary methods.

### GLP-Tox study

The NXT007 drug substance was intermittently administered with SC injections at dose levels of 0 (control), 2, 20, and 80 mg/kg (dose volume: 0.96 mL/kg) to male and female cynomolgus monkeys (5 males and 5 females per group) once every 2 weeks for 6 months to evaluate toxicity. The reversibility of toxicity was assessed during a 5-month recovery period. Systemic exposure was also assessed. Vehicle (20 mmol/L histidine-aspartate buffer containing 150 mmol/L arginine, 20 mmol/L methionine, and 0.5 mg/mL polysorbate 80, pH 6.0) was intermittently administered to animals in the control group in the same manner as the test article. During the study, toxicological observations and examinations were conducted.

### Statistical analysis

In the in vivo hemostatic study, data are presented as mean ± SE. Other data are presented as mean ± SD.

### Data and materials availability

Materials are available from Chugai Pharmaceutical Co., Ltd. under a material transfer agreement.

## RESULTS

### Identification of a lead BsAb, NXT000, with novel non-common light chains respective to the heavy chains of emicizumab

When we originally created emicizumab, a common light chain format was selected to satisfy industrial manufacturing requirements (Figure 1A).^12^ However, the use of a common light chain limited our freedom to identify the most potent light chains for the anti-FIXa arm and anti-FX arm. To overcome this limitation, we identified pairs of charge mutations in the C_H_1/C_L_ interface that could electrostatically facilitate the correct pairing of heavy and light chains,^21^ and the C_H_3/C_H_3 interface for heavy chain heterodimerization,^20^ to enable the industrial manufacturing of a non-common light chain BsAb (Figure 1B). We then screened novel light chains for each emicizumab heavy chain (anti-FIX(a) or anti-FX) using light-chain shuffled phage-displayed libraries; the common light chain of emicizumab was substituted using human naïve light chain libraries to form Fab-displayed phage libraries followed by panning against FIXa and FX (supplemental Figure 1). Based on the screening results for the in vitro cofactor activity and the binding property against each antigen, we selected NXT000 as a lead BsAb. NXT000 showed FVIIIa-mimetic cofactor activity comparable to that of emicizumab at a low concentration range in an enzymatic assay using purified coagulation factor proteins (supplemental Figure 2A).

**Figure 1.**
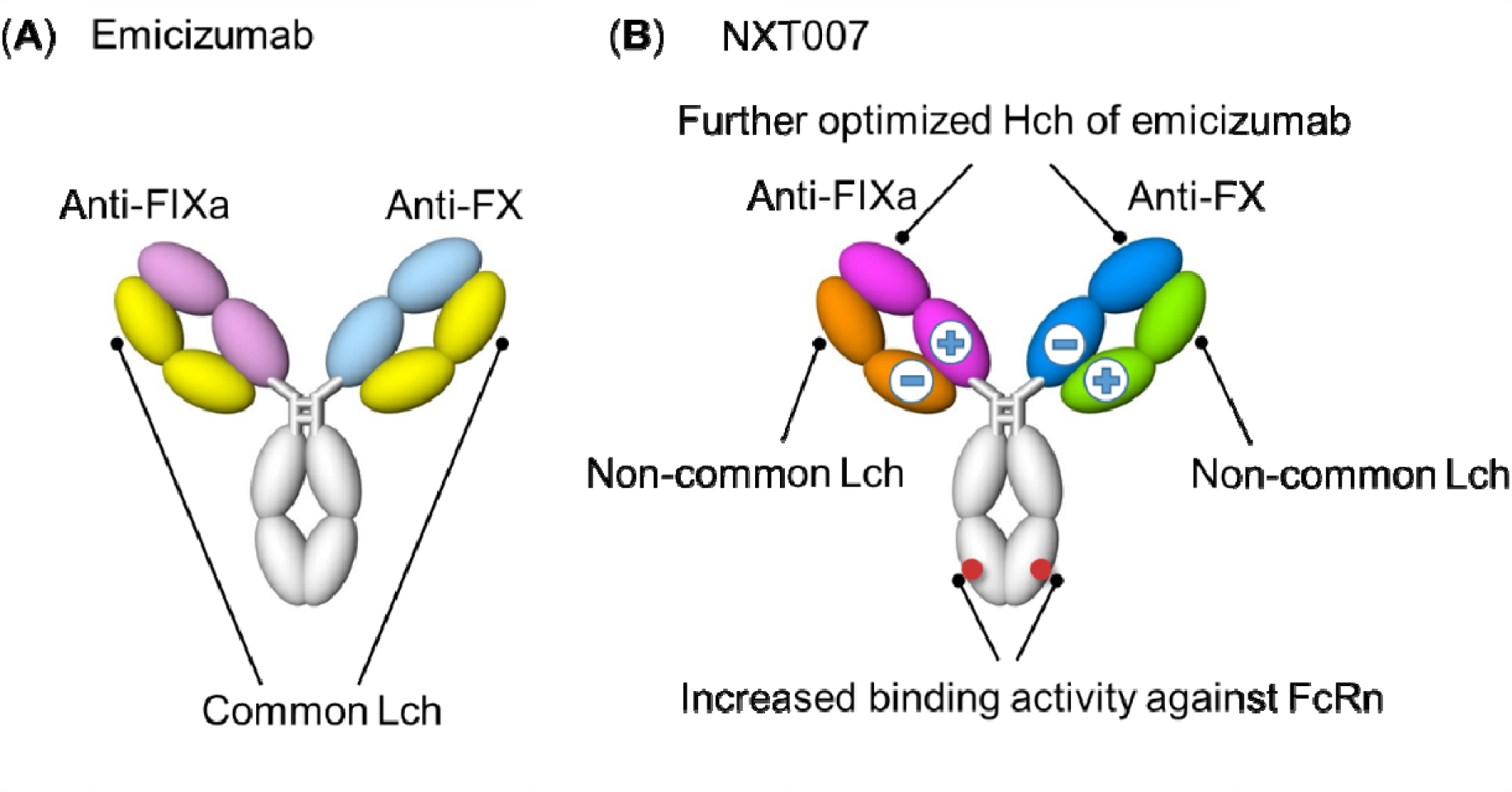
Molecular features of NXT007 and emicizumab. **(A)** Schematic illustration of emicizumab structure. The two Fab arms share a common light chain (Lch), depicted in yellow. **(B)** Schematic illustration of NXT007 structure and the introduced mutations. Optimized emicizumab-derived heavy chains and two distinct light chains from human naïve antibody libraries. Charged residue mutations are inserted into the C_H_1/C_L_ interface for correct pairing of heavy chain (Hch) and light chain (Lch).

Similar data were also observed using a plasma TG assay with activated factor XI (FXIa)-triggering condition, which was selected due to higher sensitivity than low tissue factor (TF)-triggering condition (supplemental Figure 2B). Thus, we assumed that NXT000 has the potential to achieve higher activity than emicizumab through optimization.

### Generation of NXT007 and its in vitro pharmacological profiles

To optimize NXT000, the peak height of plasma TG assays with TF-trigger was used as an indicator, since it best reflects in vivo hemostatic potential based on our retrospective analysis (supplemental Table 1; supplemental methods). After evaluating more than 5000 variants of NXT000, we successfully identified a clinical candidate named NXT007. In the enzymatic assay, NXT007 showed enhanced maximum cofactor activity at a lower concentration compared with emicizumab (Figure 2A). In FVIII-deficient patient plasma, NXT007 also showed shortened APTT (supplemental Figure 3A) and enhanced TG activity in the TF-triggering conditions (Figure 2B). In the TG assay, NXT007 showed comparable activity to emicizumab at approximately 30-fold lower concentration and showed greater activity than 100 IU/dL of recombinant human FVIII (rhFVIII) at around 120 nM (Figure 2B). Notably, the maximum TG activity of NXT007 did not exceed rhFVIII 150 IU/dL, the upper level of the non-hemophilic range. In the FXIa-triggering condition, NXT007 also showed greater activity than emicizumab. A higher concentration of NXT007 was required to show the same equivalent FVIII activity in the FXIa-triggering condition; this was also seen with emicizumab (supplemental Figure 4).

**Figure2.**
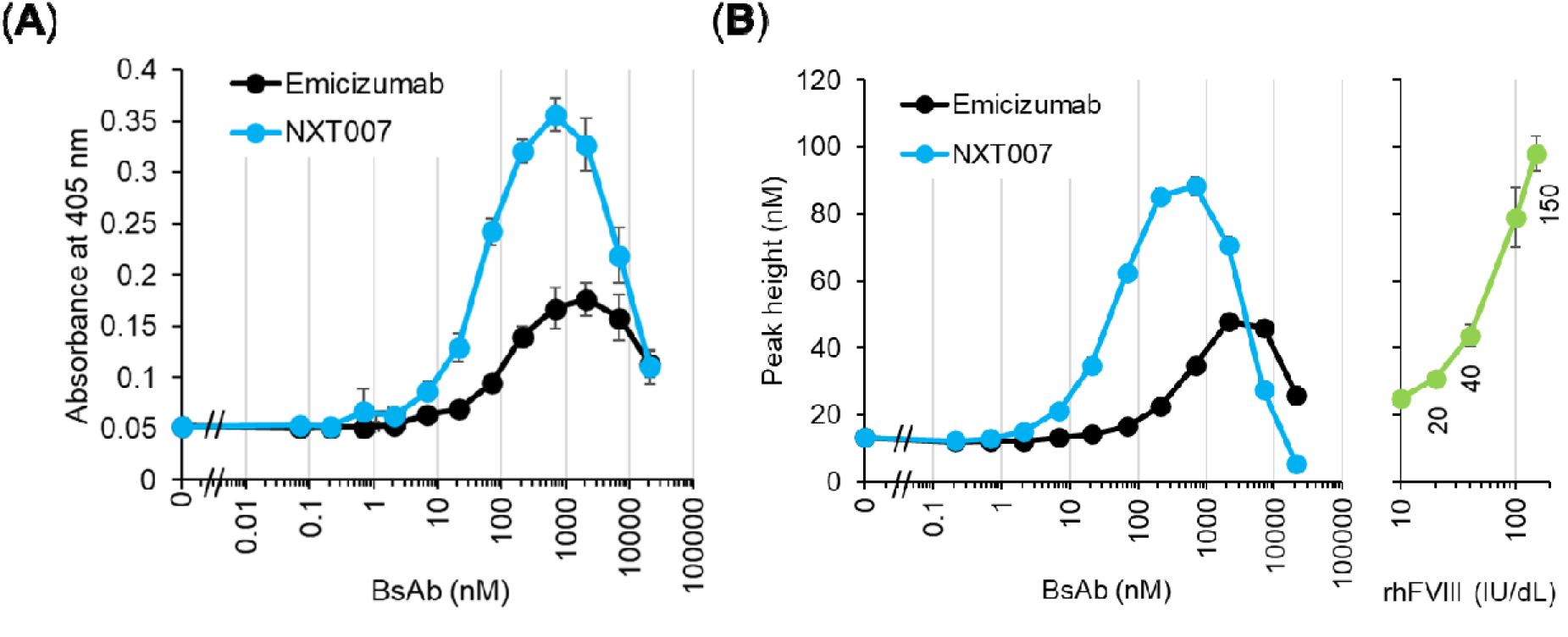
In vitro FVIIIa-mimetic cofactor activity of NXT007. **(A)** Effect of NXT007 or emicizumab on FIXa-catalyzed FX activation in an enzymatic assay using purified coagulation factors. Data are expressed as mean ± SD (n = 3) **(B)** Effect of NXT007, emicizumab or rhFVIII on thrombin generation using FVIII-deficient patient plasma. The reaction was triggered by TF. Data are expressed as mean ± SD (n = 3)

We determined that the binding affinity (*K*_D_) of NXT007 to FIX/FIXa was approximately 1 μM, which is similar to that of emicizumab.^22^ The *K*_D_ value of NXT007 to FX/FXa was approximately 30 to 40-fold reduced from emicizumab, and such enhancement of binding affinity can be explained by the improved association rate (*k*_a_) rather than dissociation rate (*k*_d_) (Figure 3A). The lower *K*_D_ value could potentially inhibit FX function since NXT007, having emicizumab’s heavy chain variants, is considered to bind to epidermal growth factor-like domain 2 (EGF2 domain) of FX as emicizumab does,^22^ which may restrict the availability of FX/FXa for the other reactions in the coagulation cascade. Thus, we carefully monitored the effect of BsAbs on FX function using PT, which is able to reflect FVIIa/TF-catalyzed FX activation and FVa/FXa-catalyzed prothrombin activation. Consequently, PT prolongation in FVIII-deficient patient plasma was minimal even in a concentration range where the TG activity of NXT007 reaches around 100 IU/dL of rhFVIII (Figure 3B; supplemental Figure 3B).

**Figure 3.**
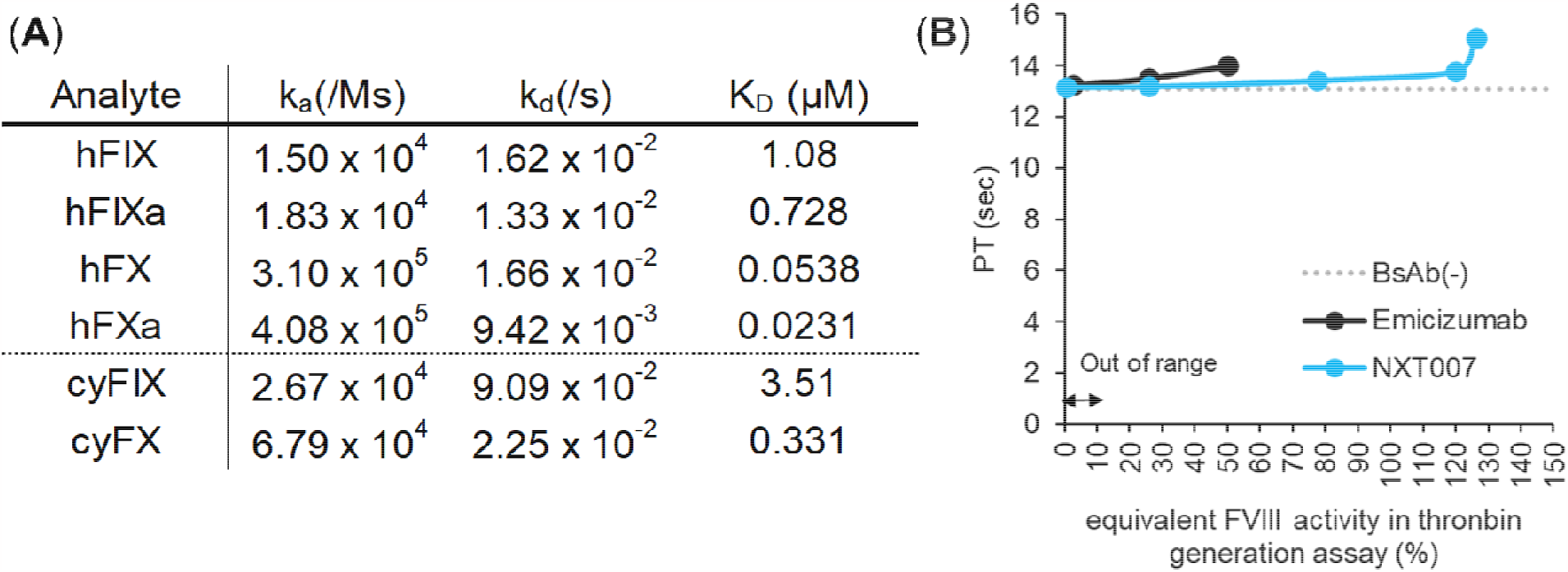
Antigen binding affinities and effect on prothrombin time of NXT007. **(A)** Antigen binding affinity of NXT007 to hFIX/hFIXa, hFX/hFXa, cyFIX and cyFX using surface plasmon resonance (SPR). The *k*_a_, *k*_d_ and *K*_D_ values are expressed as the average of three experiments. The affinity of emicizumab was previously reported.^22^ **(B)** Effect of NXT007 or emicizumab on PT using FVIII-deficient hemophilia A plasma. Equivalent-FVIII activity of NXT007 and emicizumab was calculated using the peak height of the TG assay with the TF-triggering condition calibrated by the rhFVIII standard. The analysis was carried out below the concentration range which reached maximum thrombin generation activity of NXT007 or emicizumab. Data are expressed as mean ± SD. (n = 3)

Altogether, NXT007 showed the potential to achieve a non-hemophiliac range of TG activity in FVIII-deficient plasma in vitro while having a minimal inhibitory impact on FX(a)-function in plasma.

### In vivo pharmacological profiles of NXT007

Prior to in vivo evaluation, we confirmed that NXT007 was cross reactive with cynomolgus monkey FIX/FX (Figure 3A), and dose-dependently enhanced TG activity in cynomolgus monkey plasma (supplemental Figure 5). In the experiments using monkey plasma, FXIa-triggering TG was selected due to the lack of detectability of recombinant porcine FVIII (rpoFVIII) activity, a positive control with no cross reactivity with cyVIII-2236, in the TF-triggering condition.^18^ Next, we evaluated the hemostatic activity of NXT007 using the artificially induced on-going bleeding model in acquired hemophilia A cynomolgus monkeys, which had been previously used to estimate the non-clinical equivalent-FVIII hemostatic activity of emicizumab.^18,22^ As a result, NXT007 dose-dependently suppressed bruised areas on the skin and the reduction of blood hemoglobin observed in vehicle-treated monkeys, suggesting the amelioration of bleeding symptoms (Figure 4A-B). Considering both parameters, bruised areas and hemoglobin level, at least a single IV administration of 0.075 mg/kg NXT007 exerted a level of hemostatic activity similar to twice-daily administration of recombinant porcine FVIII (rpoFVIII) 20 U/kg. At 0.075 mg/kg in the NXT007 group, the plasma concentration ranged from 1.9 μg/mL (13 nM) at Day 0 to 0.75 μg/mL (5.2 nM) at Day 3 (Figure 4C).

**Figure 4.**
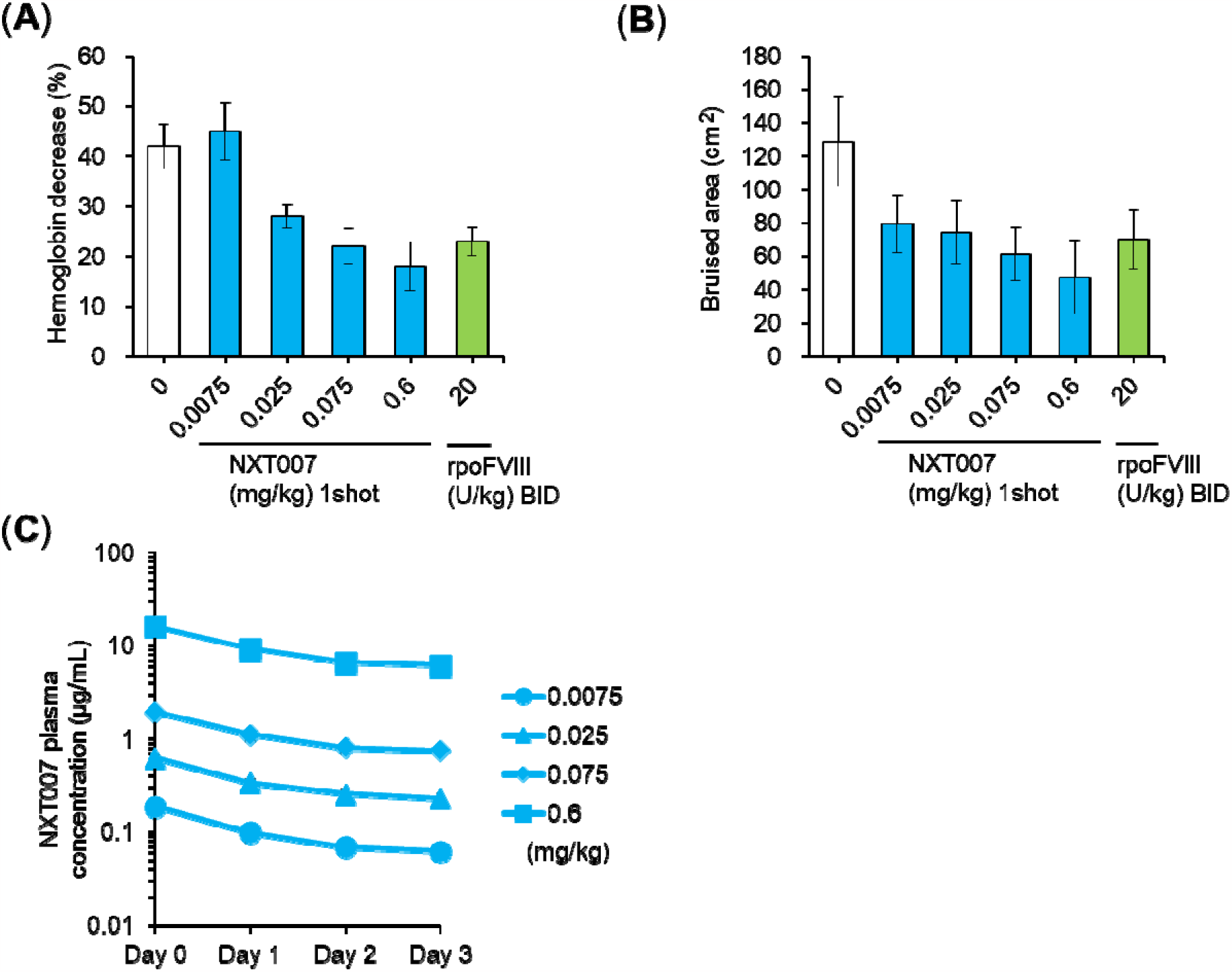
In vivo hemostatic activity of NXT007. **(A-C)** Hemostatic activity of NXT007 or rpoFVIII on acquired hemophilia A bleeding cynomolgus monkey model. Decrease of blood hemoglobin level (A) and bruised area on the skin (B) on Day 3 are shown. FVIII neutralization by cyVIII-2236 injection and bleeding induction are conducted on Day 0. Single administration for vehicles (n = 7, one animal was excluded from analysis due to lethal bleeding at Day2) and NXT007 groups (n = 6, each group), and twice daily administration for rpoFVIII groups (n = 6). Data are expressed as mean ± SE. (C) Time course of plasma NXT007 levels by ELISA. Data are expressed as mean ± SD. (n = 6)

During the experimental period, one animal in the vehicle group exhibited lethal bleeding at Day2 possibly due to the intensity of the bleeding induction method. The animal was excluded from the analysis, but this is not expected to affect the risk of overestimating NXT007 efficacy.

### In vivo pharmacokinetic study and multiple dosing simulation of NXT007

Next, the pharmacokinetics of NXT007 were assessed in a single-dose study in non-human primates (n = 4). We designed NXT007 to improve pharmacokinetics and reduce clearance by incorporating a set of mutations into the C_H_3 region to increase binding to the neonatal Fc receptor (FcRn) at acidic pH.^24^ Indeed, the clearance (CL) of NXT007 was 2.46 mL/day/kg after intravenous (IV) administration at 2 mg/kg, and CL/F in the range of 2.68–3.37 mL/day/kg after a single subcutaneous (SC) administration at 0.02, 0.2, and 2 mg/kg, which is lower than that of typical monoclonal antibodies including emicizumab,^12,18,25^ and the SC bioavailability was in the range of 82.0–92.2% (Figure 5B). The plasma elimination half-life of NXT007 was 22.1 days after IV administration at 2 mg/kg, and in the range of 19.6–24.4 days after a single SC administration at 0.02, 0.2, and 2 mg/kg (Figure 5B). With SC administration, the maximum plasma concentration of NXT007 increased in an approximately linear dose-proportional manner (Figure 5A).

**Figure 5.**
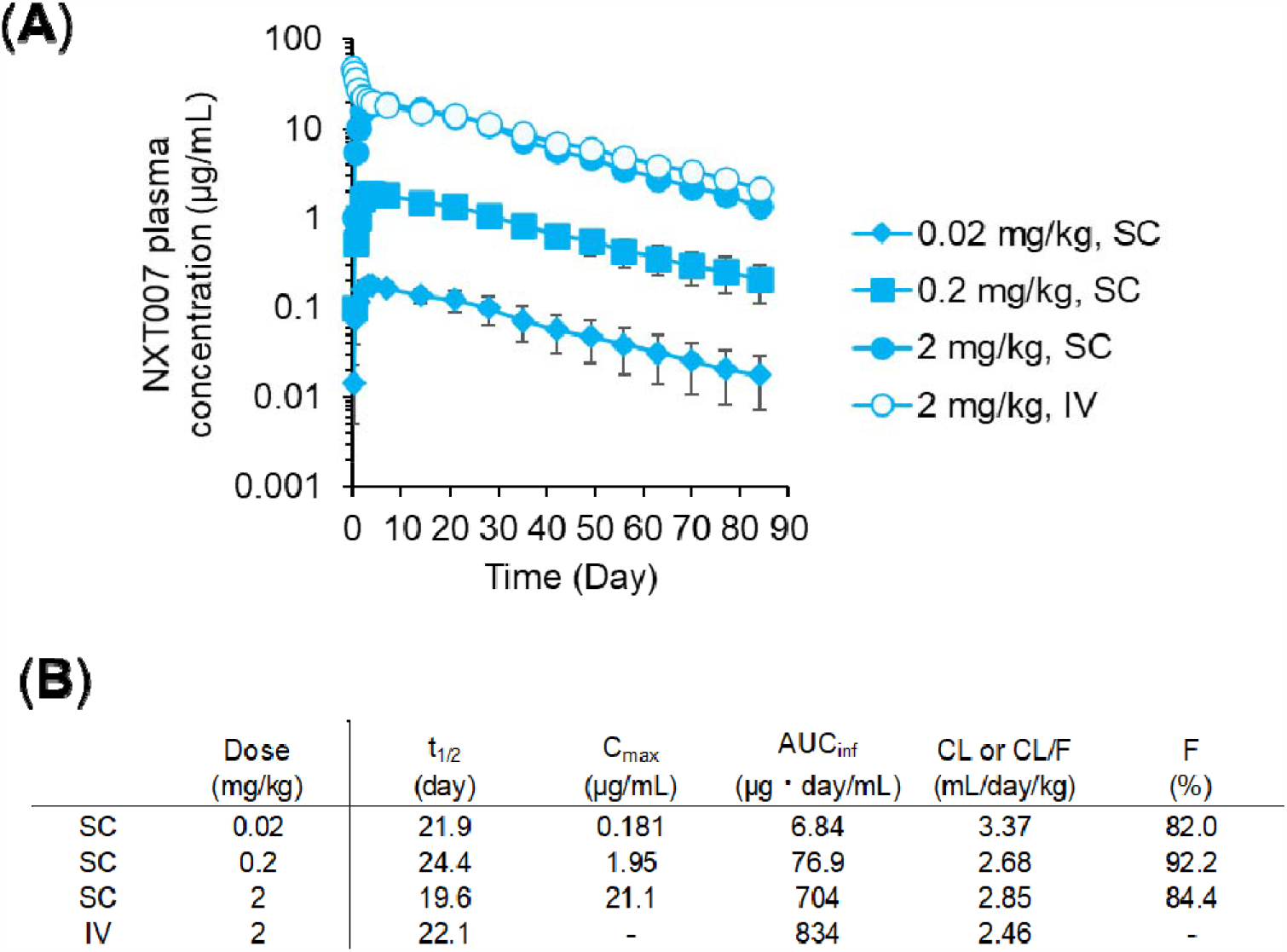
Pharmacokinetic profile of NXT007. **(A-B)** Pharmacokinetic profile of NXT007. (A) Time course of plasma concentration of NXT007 in cynomolgus monkey after 0.02-2 mg/kg intravenous (IV) or 2 mg/kg subcutaneous (SC) injection. Data are expressed as mean ± SD. (n = 4, excluded ADA-positive samples) (B) Pharmacokinetics parameters. PK parameters were calculated after excluding sampling points (n = 4, 0.02, 0.2 mg/kg, n = 2, 2 mg/kg IV and SC). –: Not applicable

### In vivo toxicology profile of NXT007

Finally, we conducted a Good Laboratory Practice (GLP) toxicology study of NXT007 (SC, every 2 weeks (Q2W), 14 doses in total at doses of 0, 2, 20, and 80 mg/kg (n = 5)). In the 6-month study in non-human primates, shortened APTT and prolonged PT were observed, which were considered to be a pharmacological effect and an exaggerated pharmacological effect. In 80 mg/kg, the prolonged PT was associated with hemorrhage-related findings in pathology.

Observed findings were considered reversible. No Observed Adverse Effect Level (NOAEL) for the 6-month GLP study in cynomolgus monkeys was 20 mg/kg SC Q2W. Detailed toxicokinetic parameters are shown in supplemental Table 2.

## DISCUSSION

It is now established with both non-clinical data and subsequent clinical experience that emicizumab converts a PwHA to the mild hemophilia range of FVIII:C equivalence. Our goal was to develop a BsAb, reaching the levels of 100 IU/dL of FVIII (equivalent to standard normal plasma) in the in vitro TF-trigger TG assay to further improve the steady state level of prophylaxis protection near to the normal hemostatic range (40–150 IU/dL of FVIII-equivalence), thus theoretically preventing breakthrough bleeding as well as joint bleeds. We hypothesized that using a four-chain BsAb format instead of emicizumab’s common light chain format would offer greater engineering flexibility and result in higher activity. Additionally, we aimed to maintain the clinically validated FIX(a) and FX epitopes used by emicizumab. Thus, we utilized phage libraries, which display Fabs consisting of emicizumab’s heavy chains and human naïve light chain libraries. We identified two distinct novel light chains and through extensive optimization of the Fab domains ultimately developed NXT007, which achieved a maximum activity of over 100 IU/dL of FVIII in the TF-trigger TG assay.

During our optimization process, we aimed to balance the conflicting objectives of increasing the formation of FIX(a)–BsAb–FX ternary complex while minimizing any inhibitory effect on the innate functions of FIX(a) and/or FX as much as possible. To accomplish this, we employed a strategy to keep the *K*_D_ of the anti-FIX(a) arm similar to that of emicizumab (micro molar range) while strengthening the *K*_D_ of the anti-FX arm by increasing *k*_a_ rather than decreasing *k*_d_. Our hypothesis was that faster association and dissociation to FX minimizes the coagulation inhibitory effect of the anti-FX arm while maintaining a high turnover rate to form FIX(a)– NXT007–FX ternary complex. We were able to examine the inhibitory effect of the anti-FX arm using the in vitro PT assay. Indeed, this assay suggested that NXT007 did not prolong PT as much as emicizumab when their equivalent-FVIII TG activity was the same (Figure 3B), expecting to have a wide therapeutic window like emicizumab.

After confirming the in vitro profiles of NXT007, we conducted in vivo studies to confirm whether, like emicizumab, the in vitro potency reflected the in vivo hemostatic potential. However, due to the ethical limitations on the use of non-human primates, a direct comparison between NXT007 and emicizumab in vivo could not be made. Thus, the results of this NXT007 study are discussed in relation to those of the previous emicizumab study (supplemental Figure 6). In both studies, the mean values of hemoglobin decrease in the control groups were comparable. Additionally, hemostatic activity in the rpoFVIII groups was similar, despite the different doses of rpoFVIII used; 20 or 10 U/kg twice daily in this study or the previous study, respectively (supplemental Figure 6). We think these two studies can be compared because both dosing regimens were set to achieve a substantial level of hemostatic activity and keep the disease severity mild. These studies demonstrated that a single IV dose of NXT007 as low as 0.075 mg/kg had a significant hemostatic effect similar to that of twice-daily doses of 20 U/kg rpoFVIII (Figure 4A-B), and that a single IV dose of 3 mg/kg emicizumab had a hemostatic effect roughly comparable to twice-daily doses of 10 U/kg rpoFVIII.^18^ The mean plasma concentration of NXT007 and emicizumab just after the single IV administration at the above doses was 1.9 μg/mL (Figure 4D) and 61 μg/mL, respectively.^18^ The difference in the plasma BsAb concentrations was approximately 30-fold. These in vivo data are roughly consistent with the results of in vitro TG assays for NXT007 and emicizumab, as NXT007 showed an approximately 30-fold lower dose shift than emicizumab in the in vitro TG assay (Figure 2B). In other words, the predictability of emicizumab’s in vivo hemostatic activity using in vitro TG assays (supplemental Table 1) was reproduced for NXT007. However, this conclusion should be interpreted with caution as the number of animals used in these studies was relatively small and the environmental differences between the NXT007 and emicizumab studies may have had a potential impact.

A single-dose pharmacokinetic study using cynomolgus monkeys demonstrated that the pharmacokinetic profile of NXT007 was comparable or slightly better than that of emicizumab (Figure 5). This is likely due to the benefit of increased FcRn binding at acidic pH introduced by Fc engineering.^24^ The prediction of human pharmacokinetics for such Fc engineered antibodies is still challenging because very few have been tested in humans. However, motavizumab-YTE, another type of Fc engineered antibody with increased FcRn binding at acidic pH showing a similar clearance in cynomolgus monkeys to NXT007,^26^ demonstrated a half-life of 70–100 days in humans. ^27^ If this type of Fc engineering generally led to a large increment of half-life in human settings, we might expect NXT007 also to show such a long half-life in clinic.

We also simulated a possible subcutaneous dosing regimen of NXT007 using non-clinical efficacy and cynomolgus pharmacokinetic data. We expectedly assume that authentic in vitro TG activity of NXT007 might be reflected to hemostatic activity in PwHA when extrapolating with results of emicizumab. This is because in vitro TG activity of emicizumab was roughly consistent with in vivo hemostatic activity as above mentioned. Furthermore, the results of in vivo study were also roughly concordant with hemostatic potential in PwHA in clinic.^15,28^ Thus, together with the cynomolgus pharmacokinetic data, NXT007 is expected to maintain a non-hemophilia range (>40 IU/dL of rhFVIII) in PwHA by longer dosing intervals such as every 4 weeks (Q4W) SC administration (Supplemental Figure 7). Moreover, the effect on PT prolongation would be minimal at this range given that 40-100 IU/dL of rhFVIII in TG can be achieved by 4.4-22.3 μg/mL of NXT007 (Figure 3B, Supplemental Figure 3).

In the GLP-tox study, at Q2W SC dosing of 20 mg/kg NXT007, PT prolongation was observed without any pathological changes, while at 80 mg/kg, PT prolongation was accompanied with hemorrhage-related findings probably caused by FX inhibition due to the high concentrations of NXT007. However, the C_max_ achieved in the 20 mg/kg SC Q2W study (approximately 1000 μg/mL as shown in Supplemental Table 2) was much higher than the NXT007 concentration that could maintain non-hemophiliac range; more than 200-fold higher than the concentration that exerted 40 IU/dL (4.4 μg/mL) and more than 40-fold higher than the concentration that exerted 100 IU/dL of rhFVIII in the TF-triggering condition (22.3 μg/mL). Irrespective of the degree of hemorrhagic observation in the GLP-tox study, considerable care should be taken in PT-prolongation in the real-world settings. The GLP-tox study was performed in non-hemophiliac animals with neither thrombotic nor hemorrhagic invasion.

In conclusion, our non-clinical findings on the efficacy, safety, and pharmacokinetics of NXT007 indicate that it will be able to safely maintain a non-hemophiliac range of coagulation potential in PwHA. Evaluations of safety, tolerability, pharmacokinetics, pharmacodynamics, and efficacy of NXT007 in PwHA are ongoing.

## Acknowledgments

We thank the donors and patients who consented to the use of their cells for the studies. We also thank colleagues in Chugai Pharmaceutical Co., Ltd. and Chugai Research Institute for Medical Science Inc; S. Naoi, S. Nagatomo, Y. Wakahara, K. Komura, Y. Kaneta, S. Komatsu, B. Juan, Y. Ikeda, N. Sato, M. Uematsu, A. Ohba, Y. Oyama, A. Maeno, K. Matsunaga, S. Hiranabe, T. Kato for antibody preparation and characterization; S. Kadono, T. Torizawa, J. Okude for structural analysis; F. Isomura, Y. Ochiai for in vitro experiments; R. Saito, M. Namba, T. Hasegawa, M. Fukazawa, H. Azabu, M. Hiranuma, H. Arakawa, R. Takemoto, Y. Miura, T. Koike, T. Nakatogawa, H. Tai for in vivo experiments.

## Author contributions

Y.T.-I., T.S., H. Koga, Kazuki Y., H. Sanada, N.S., A. Matsusaki, and T.K. wrote the manuscript; Y.T.-I, H. Koga, K.K., K.E., M.F., M.I., Y.I., M.O., M.S., and T.W. designed, produced, and characterized the antibodies; T.S., Kazuki Y., E.J., A. Muto, and K.N. executed the pharmacological studies and analyzed the assay data; K.A., S.I., S.i.-K., H. Sanada, N.S., and Koji Y. executed the PK and toxicological studies and analyzed the assay data; Z.S., H. Shiraiwa, H. Konishi, Y.K., K.H., T.K., and T.I. designed and supervised the studies.

