## Supplemental figures and tables for "A bispecific antibody NXT007 exerts a hemostatic activity in hemophilia A monkeys enough to keep a non-hemophiliac state"

**Supplemental Methods**

**Materials**

Emicizumab (ACE910), anti-primate FVIII neutralizing antibody (cyVIII-2236), and B domain-deleted recombinant porcine FVIII (rpoFVIII) were prepared as previously described.^1^

**Determination of a screening condition for FVIII-function mimetic activity of emicizumab variants**

Describing the hemostatic activity of emicizumab from in vitro assay is challenging due to its unique mode of action, unlike FVIII, which can be predictably described based on international units. We re-examined various in vitro studies of emicizumab to determine the most appropriate parameter to reflect its in vivo hemostatic potential. By analyzing a series of in vitro datasets on emicizumab at 50 μg/mL,^2-5^ we found that the conversion factor determined by the peak height of plasma TG assays with low tissue factor-trigger (TF-trigger) was 0.28, similar to the conversion factor (around 0.3) obtained from previous non-clinical in vivo studies.^2,6^ Peak height, defined as the highest thrombin concentration in the TG assay, can reliably indicate hyper- or hypo-coagulability^7^ and showed concentration-dependency of recombinant human FVIII at 10-100 IU/dL, making it a suitable parameter to screen and evaluate emicizumab variants targeting 100 IU/dL activity.

**Identification of novel non-common light chains from phage library**

The light chain shuffled Fab-displayed phage libraries were constructed in Chugai Pharmaceutical Co., Ltd. as follows: Human naïve kappa or lambda gene libraries derived from 43 healthy volunteers were combined with genes of either Q499 or J327 (both are the heavy chains of emicizumab), and then inserted into a phagemid vector to display Fab domains on phage. Phages were produced using E coli, followed by bio-panning using biotin-labelled human FIXa or FX as antigens. After several rounds of panning, phage ELISA was performed to check the binding against each antigen and select the pools which were then used for conversion of IgG expressible by mammalian cells such as Expi293 (ThermoFisher Scientific). Approximately 1000 IgGs were produced for each anti-FIX(a) or anti-FX antibody screening.

**Generation of the efficacy enhanced anti-FIXa/FX bispecific antibody NXT007**

NXT007 was generated by optimizing NXT000. We first aimed to improve the maximum FVIIIa-mimetic cofactor activity, especially the maximum equivalent-FVIII TG activity, so that it would be much more potent than emicizumab. To do this, we used the COSMO (Comprehensive substitution for multidimensional optimization) approach,^8^ each amino acid residue in all complementarity determining regions (CDRs) and in some positions in the framework regions of NXT000’s heavy and light chains were replaced with a different natural amino acid except cysteine on a one-by-one basis. We selected mutations to combine which were able to increase cofactor activity in the enzymatic assay compared to NXT000. Then, the effective combinations were further assessed using TG assays. We also introduced other mutations to achieve an isoelectric point (pI) difference between the two arms for purification processes, to decrease non-specific binding, to improve solubility and thermal stability, and to minimize immunogenicity in the same manner as emicizumab.^9,10^ Consequently, we identified a clinical candidate which we named NXT007.

**Binding kinetics analysis by surface plasmon resonance (SPR)**

Each antibody was captured onto Sure Protein A (MabSelect SuRe) immobilized on a CM4 sensor chip (Cytiva), after which hFIX, hFIXa, hFX, hFXa (Enzyme Research Laboratories, Inc.), cyFIX, or cyFX (prepared in-house) were injected over the flow cells. To suppress the non-specific interaction of the analyte with the sensor chip, the surface negative charge of the sensor chip was reduced by the pre-treatment, described as follows. First, carboxyl groups on Flow Cell 1 (FC1) and Flow Cell 2 (FC2) were activated by injecting a mixed solution of equal volumes of NHS and EDC. Then, the active groups were blocked by injecting ethanolamine-HCl. The sensor chip surface was regenerated using 25mM NaOH. Kinetic parameters were determined by fitting the sensorgrams obtained by subtracting the FC1 sensorgram from the FC2 sensorgram with 1:1 binding model using Biacore T200 evaluation software, version 2.0 (Cytiva). The average values of kinetic parameters were calculated from three measurements.

**Enzymatic assay for FIXa-catalyzed FX activation**

The assay system consisted of 1 nM human FIXa (Enzyme Research Laboratories), 140 nM human FX (Enzyme Research Laboratories), 4 μM phospholipid (10% phosphatidylserine, 60% phosphatidyl-choline, and 30% phosphatidylethanolamine), and bispecific antibodies. FXa generation was measured at room temperature for 1 min in TBS containing 1 mM CaCl_2_, and 0.1% (wt/vol) BSA (pH 7.6). We stopped the reaction by adding EDTA. After adding S-2222 chromogenic substrate (Cromogenix), we measured absorbance at 405 nm to determine the rate of FXa generation. Data were collected in triplicate.

**Thrombin generation (TG) assays**

For the evaluation of activity in cynomolgus monkey plasma, pooled citrated plasma of 6 cynomolgus male monkeys which contains 300 μg/mL of cyFVIII-2236, and BsAb or rpoFVIII was used for plasma sample and intrinsic triggering solution consisting of 310 pM human FXIa and 20 μM synthetic phospholipid was used. For calibration, 20 µL of Thrombin Calibrator (Thrombinoscope BV) was added instead of the triggering solution. To initiate the reaction, 20 µL of FluCa reagent prepared from FluCa kit (Thrombinoscope BV) was dispensed by the instrument as programmed. We analyzed the thrombograms and peak height using the instrument’s software. Data were collected in triplicate.

**Activated partial thromboplastin time (APTT) and prothrombin time (PT)**

We used FVIII-deficient human plasma (George King Bio-Medical) for APTT and PT assay, which were treated with each concentration of BsAb or rhFVIII (Bayer AG). Data were collected in triplicate. For the analysis of the correlation of equivalent-FVIII activity of NXT007 or emicizumab and PT value, equivalent-FVIII activity was calculated using the peak height of the TG assay with the TF triggering condition calibrated by the rhFVIII standard. The analysis was carried out below the concentration range which reached maximum thrombin generation activity of NXT007 or emicizumab.

**In vivo hemostatic study in an acquired hemophilia A model**

On Day 0, to establish an acquired hemophilia A state, the animals received an intravenous injection of cyVIII-2236 (10 mg/kg). Two hours thereafter, the animals were anesthetized by isoflurane inhalation. Then, the following 2 surgical procedures were performed: 1 cm-deep insertion of an 18G-needle into the muscle at 22 sites (2 sites in each upper arm, 3 sites in each forearm, 4 sites in each inside thigh, and 2 sites in each outside thigh), and subcutaneous exfoliation by 3 cm-long insertion of a tip of forceps beneath the abdominal skin at 2 sites. After administration of buprenorphine, an analgesic drug, the animals were recovered from the anesthesia. (The animals also received this analgesic treatment on the following days.) Although bleeding symptoms were not apparent at that time, visible hematoma at the sites of subcutaneous exfoliation became apparent about 6 to 8 hours later. The animals then received intravenous administration of NXT007 (0 (Vehicle), 0.0075, 0.025, 0.075 or 0.6 mg/kg; n = 7 for vehicle, and n = 6 for each NXT007 group), rpoFVIII (20 U /kg; n = 6). To the animals in the rpoFVIII group, rpoFVIII was intravenously administered once on Day 0 and twice daily on Days 1 and 2 (in the morning and evening; a total of 5 administrations). In the morning on Days 1, 2 and 3, the animals were anesthetized to measure the bruised areas. After the evaluation on Day 3, the animals were killed humanely. Citrated blood was collected before and 2 hours after the cyVIII-2236 injection, about 10 min after the test item administration on Day 0, and before measuring the bruised area on Days 1, 2, and 3. The decrease in blood hemoglobin level was expressed as a percentage on Day 0 (before initial NXT007 or rpoFVIII injection).

**Pharmacokinetic study and simulation**

The concentration of NXT007 in monkey plasma was measured by ELISA assay. An anti-NXT007 rabbit IgG was dispensed onto a microplate and incubated for 1 hour at room temperature. After each plasma sample was added to the plates and removed, another anti-NXT007 mouse IgG was reacted with samples. Anti-mouse IgG with HRP label was used as detection antibody. The signal was detected by micro plate reader. Biotin labeled NXT007 and digoxigenin labeled NXT007 were sandwiched to detect anti-NXT007 antibodies. The signal was detected by micro plate reader in PK and ADA assays.

The pharmacokinetic parameters were analyzed by non-compartmental analysis with Phoenix Winnonlin (version 6.4, Certara LP) excluding ADA positive animals or sampling points. Multiple-dosing simulations were performed with SAAM II version 1.2 (SAAM Institute, Seattle, WA, USA) based on a 2-compartment model with first order absorption kinetics.

**Supplemental Tables**

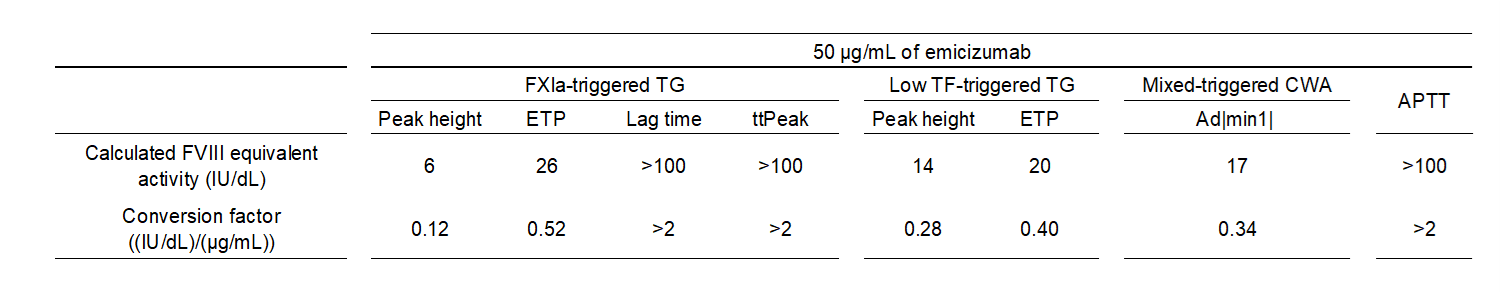

**Supplemental Table 1. Conversion factor of emicizumab to FVIII equivalent activity**

Conversion factors (coefficient) from emicizumab (μg/mL) to FVIII-equivalent activity (U/dL) were calculated based on reported in vitro data including FXIa-triggered TG assay, low TF-triggered TG assay, Mixed-triggered CWA and APTT.^2-5^

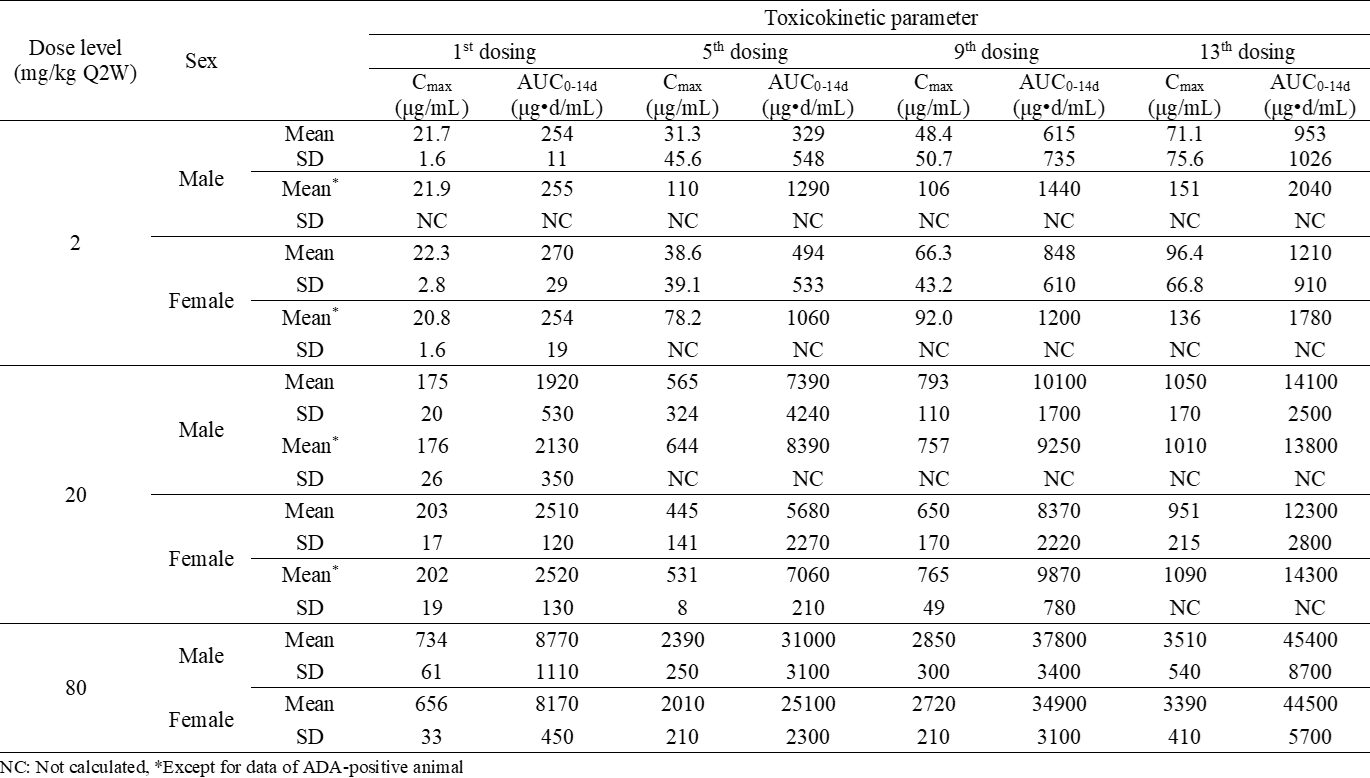

**Supplemental Table 2. Toxicokinetic parameters after administration of NXT007 in cynomolgus monkeys**

NXT007 was subcutaneously administered once every 2 weeks, 14 doses in total at doses of 0, 2, 20, and 80 mg/kg (n = 5). NC: Not calculated, *Except for data of ADA-positive animals.

**Supplemental Figures**

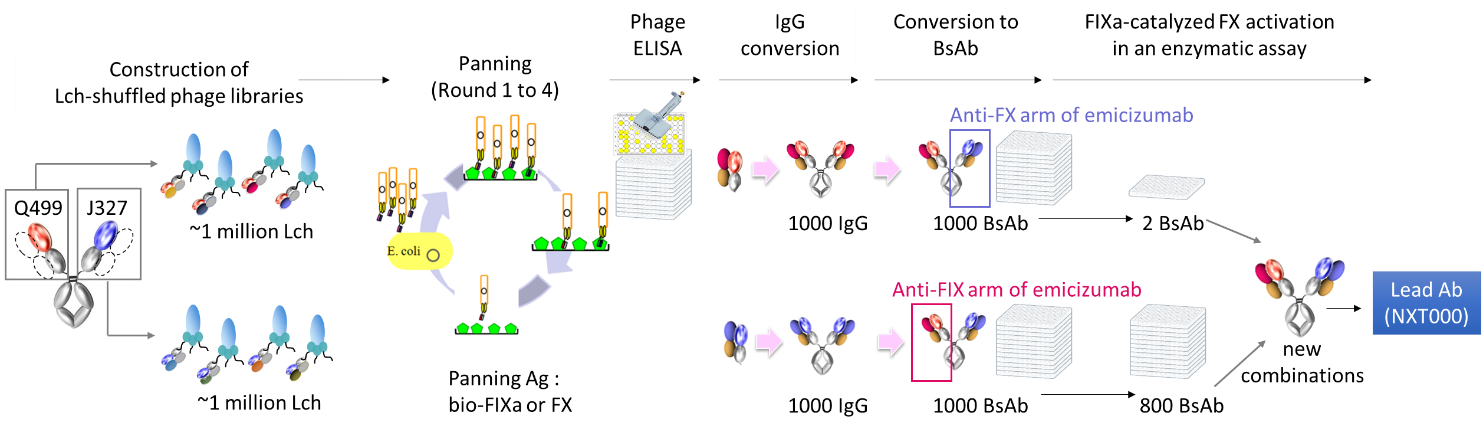

**Supplemental Figure 1. Flow to identify NXT000**

Human naïve kappa or naïve lambda gene libraries were combined with the heavy chain of emicizumab (either Q499 or J327) then Fab-displayed M13 phages were produced. Several rounds of panning were conducted using biotin-labelled human FIXa or FX. After phage ELISA (enzyme-linked immunosorbent assay) to check binding to FIXa or FX, Fabs were converted into IgGs. Approximately 1000 IgGs were produced for each anti-FIXa and anti-FX antibody, combined with either arm of emicizumab to create BsAb, and screened by FIXa-catalyzed FX activation in an enzymatic assay to identify the lead Ab, NXT000.

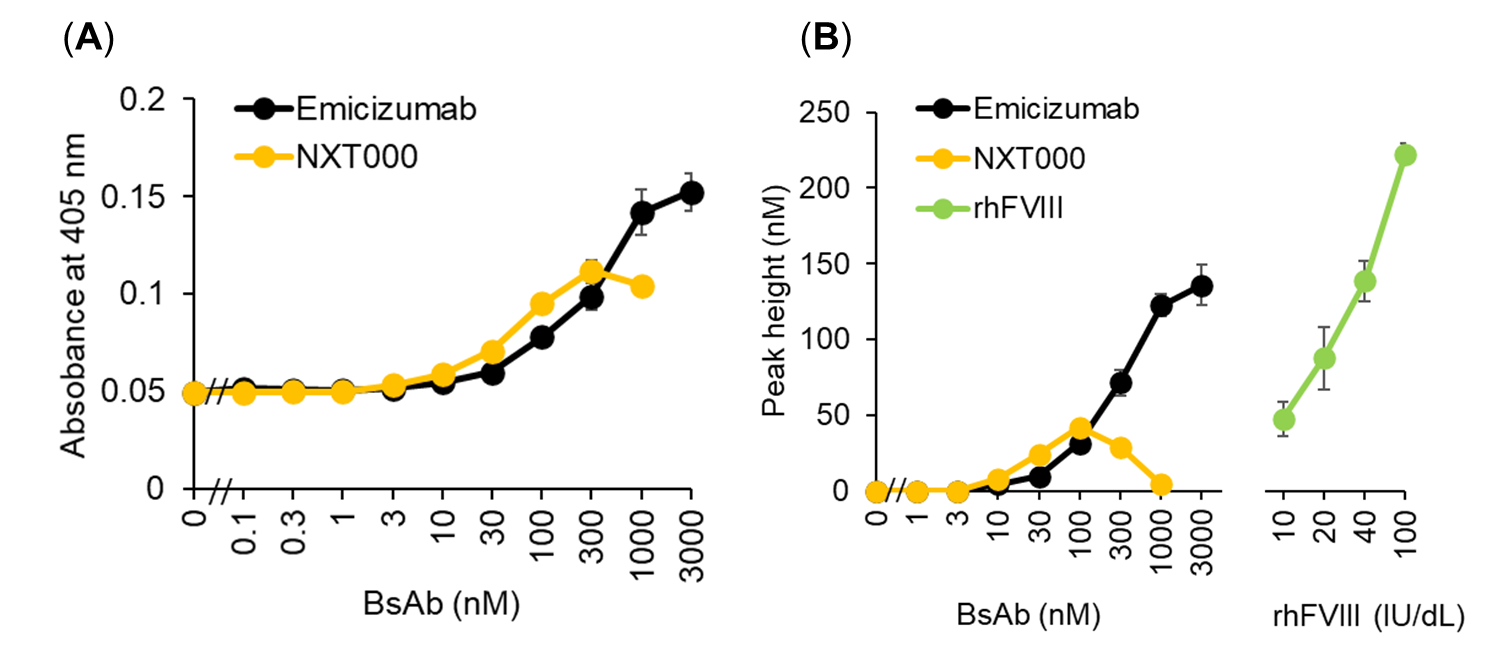

**Supplemental Figure 2. In vitro FVIIIa-mimetic cofactor activity of NXT000**

(A) Effect of NXT000 or emicizumab on FIXa-catalyzed FX activation in an enzymatic assay using purified coagulation factors. (B) Effect of NXT000, emicizumab or rhFVIII on thrombin generation using FVIII-deficient patient plasma. The reaction was triggered by FXIa. Data are expressed as mean ± SD (n = 3).

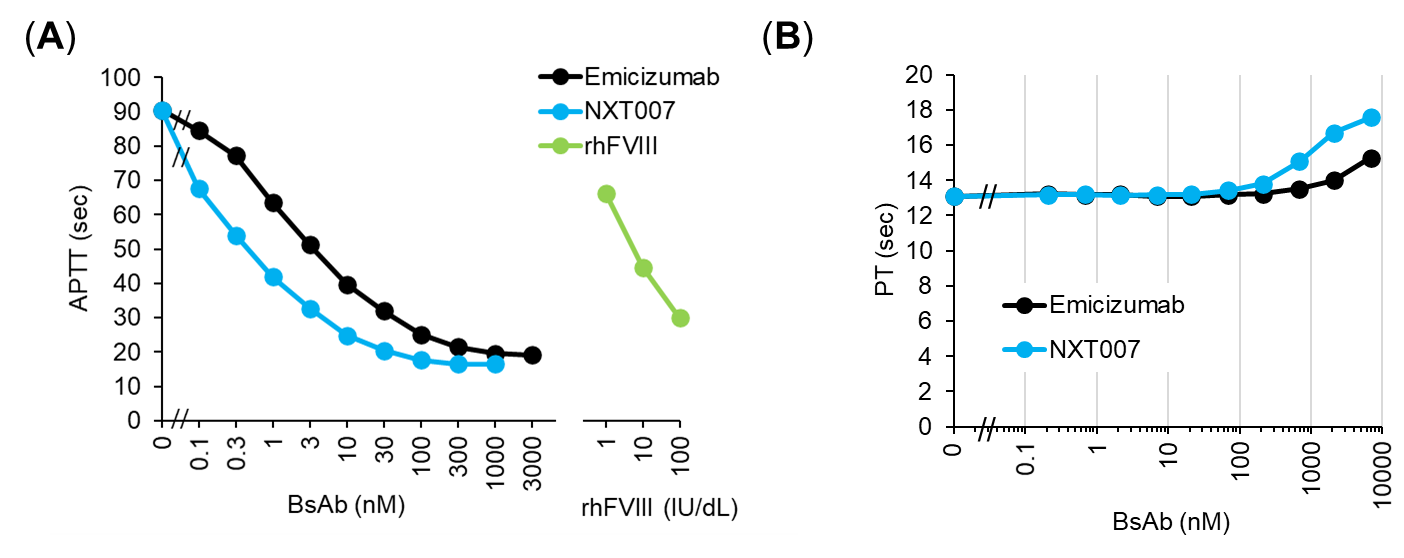

**Supplemental Figure 3. Clotting activity of NXT007 treated plasma**

(A) Effect of NXT007, emicizumab or rhFVIII on APTT in FVIII deficient patient plasma. (B) Effect of NXT007 on PT in FVIII-deficient patient plasma. Data are expressed as mean ± SD (n = 3). At higher concentrations (>69 nM) of NXT007, PT was prolonged, presumably due to increased FX(a)-occupancy, which had also been observed with emicizumab.

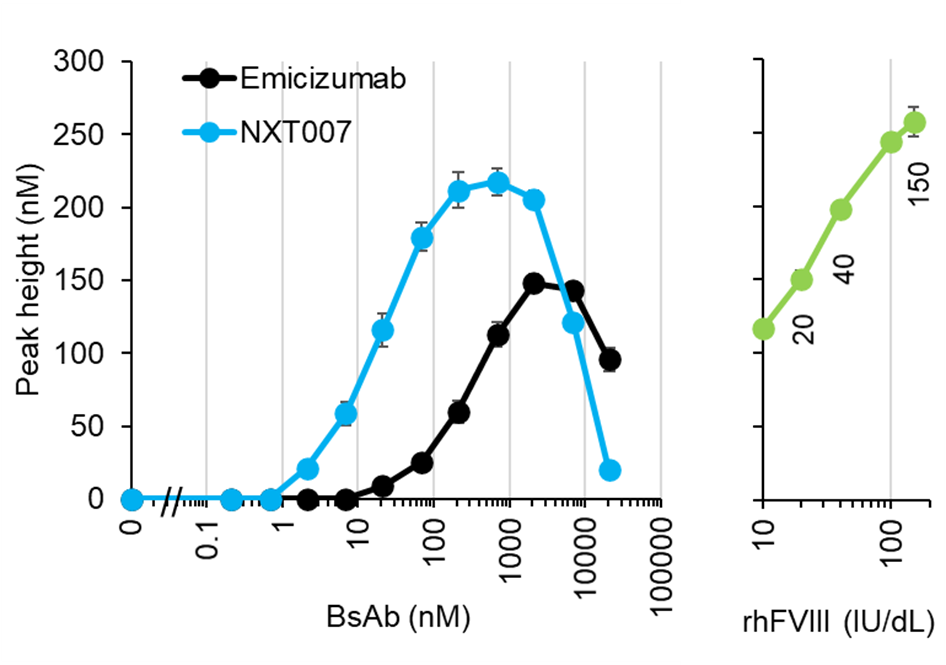

**Supplemental Figure 4. In vitro FVIIIa-mimetic cofactor activity of NXT007**

Effect of NXT007, emicizumab or rhFVIII on thrombin generation using FVIII-deficient patient plasma. The reaction was triggered by FXIa. Data are expressed as mean ± SD (n = 3).

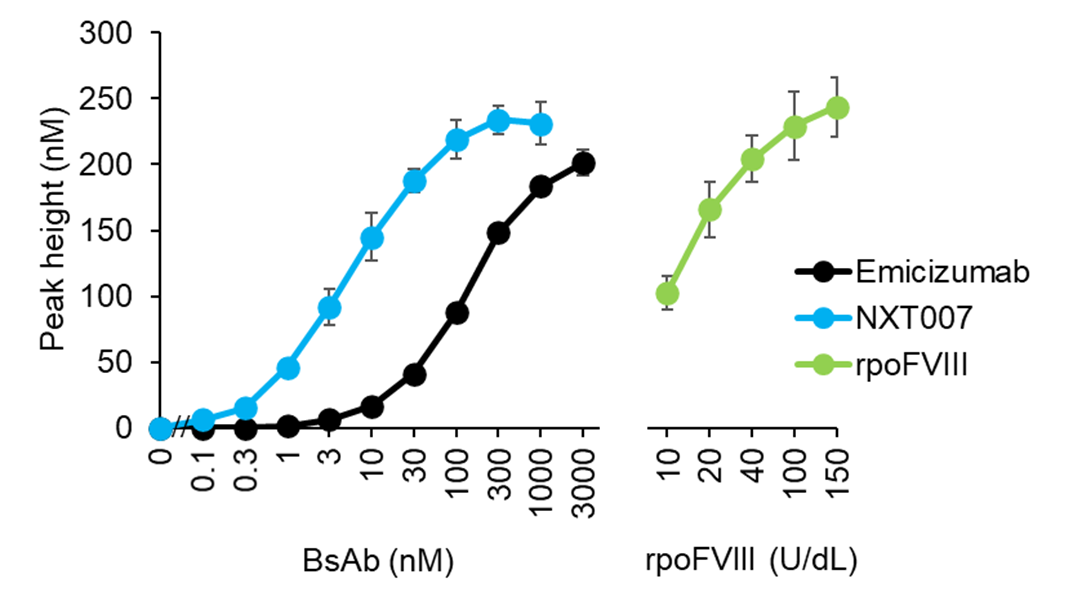

**Supplemental Figure 5. Thrombin generation activity of NXT007 in cynomolgus monkey plasma**

Effect of NXT007, emicizumab or rpoFVIII on thrombin generation using FVIII-neutralized cynomolgus monkey plasma. The reaction was triggered by FXIa. Data are expressed as mean ± SD (n = 3).

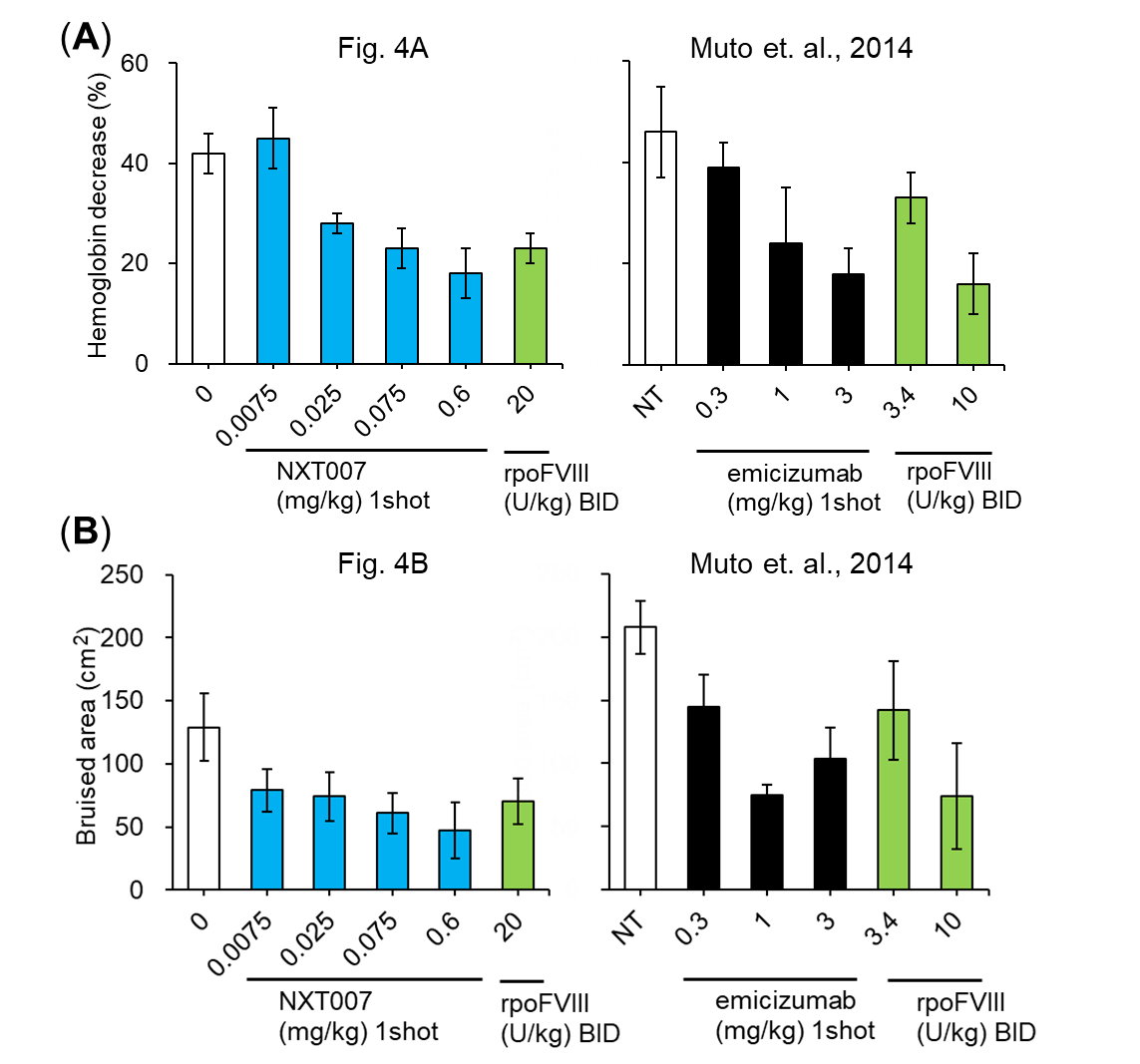

**Supplemental Figure 6. Historical comparison of in vivo hemostatic activity of NXT007 and emicizumab**

(A-B) Hemostatic activity of NXT007 and rpoFVIII in an acquired hemophilia A bleeding cynomolgus monkey model in the current study, and of emicizumab and rpoFVIII in a previous report.^2^ Blood hemoglobin level (A), Bruised area on the skin (B) on Day 3 are shown. FVIII neutralization by cyVIII-2236 injection and bleeding induction were conducted on Day 0. Single administration for vehicle (n = 7, one animal was excluded from analysis due to lethal bleeding at Day2) and NXT007 groups (n = 6, each group), and twice daily administration for rpoFVIII groups (n = 6) in the current study. No drug treatment (NT, n = 6) and single administration for emicizumab (n = 4, each group), and twice daily administration for rpoFVIII groups (n = 4, each group) in the previous study. Data are expressed as mean ± SE.

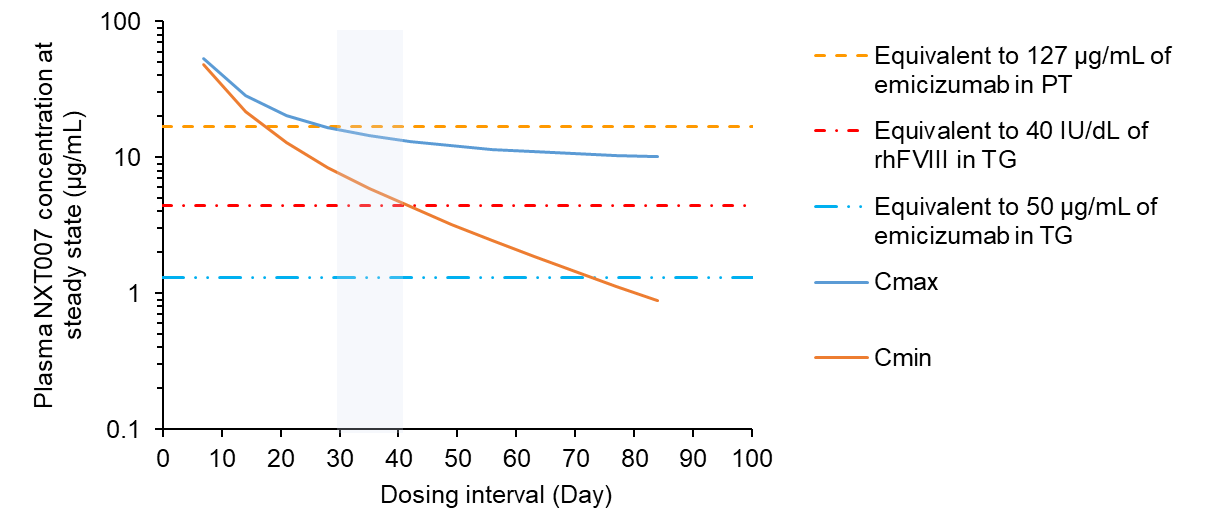

**Supplemental Figure 7. Pharmacokinetic simulation of NXT007**

Simulation of plasma concentration of NXT007 using cynomolgus monkey pharmacokinetics data. Orange curve and blue curve represent predicted plasma trough and maximum NXT007 concentration by 1.0 mg/kg, SC administration, respectively. As a reference, lines representing NXT007 concentration equivalent to 40 IU/dL of rhFVIII in TG, 50 μg/mL of emicizumab, and 127 μg/mL of emicizumab in PT are shown. Emicizumab was safely administered to patients for several years with the highest concentrations in blood reaching 872 nM (127 μg/mL), which was calculated to be highest average steady-state peak value based on the average C_min_ and the ratio of C_max_/C_min_ observed in the clinical studies of emicizumab.^11,12^
